# Functional diversity of brain networks supports consciousness and verbal intelligence

**DOI:** 10.1101/336859

**Authors:** Lorina Naci, Amelie Haugg, Alex MacDonald, Mimma Anello, Evan Houldin, Shakib Naqshbandi, Laura E. Gonzalez-Lara, Miguel Arango, Christopher Harle, Rhodri Cusack, Adrian M. Owen

**Affiliations:** Trinity College Institute of Neuroscience, Trinity College Dublin, Dublin, Ireland; Department of Psychiatry, Psychotherapy and Psychosomatics, Psychiatric Hospital, University of Zurich, Zurich, Switzerland; Faculty of Medicine, University of Toronto, Toronto, Canada; Schulich School of Medicine & Dentistry, Western University, London, Canada; Brain and Mind Institute, Western University, London, Canada; Department of Anesthesia and Preoperative Medicine, Schulich School of Medicine & Dentistry, Western University, London, Canada

## Abstract

How are the myriad stimuli arriving at our senses transformed into conscious thought? To address this question, in a series of studies, we asked whether a common mechanism underlies loss of information processing in unconscious states across different conditions, which could shed light on the brain mechanisms of conscious cognition. With a novel approach, we brought together for the first time, data from the same paradigm—a highly engaging auditory-only narrative—in three independent domains: anesthesia-induced unconsciousness, unconsciousness after brain injury, and individual differences in intellectual abilities during conscious cognition. During external stimulation in the unconscious state, the functional differentiation between the auditory and fronto-parietal systems decreased significantly relatively to the conscious state. Conversely, we found that stronger functional differentiation between these systems in response to external stimulation predicted higher intellectual abilities during conscious cognition, in particular higher verbal acuity scores in independent cognitive testing battery. These convergent findings suggest that the responsivity of sensory and higher-order brain systems to external stimulation, especially through the diversification of their functional responses is an essential feature of conscious cognition and verbal intelligence.

## Introduction

Understanding the brain mechanisms of conscious cognition is one of the great frontiers of cognitive neuroscience. A much-researched yet unresolved question is how the myriad sensory inputs arriving at our senses become integrated into meaningful representations that inform cognitive performance and give rise to individual differences in intellectual abilities. In the conscious brain, cognition is thought to arise from iterative interactions among brain regions of graded functional specialization. These include sensory-driven, e.g., auditory and visual, regions on one end of the functional hierarchy, and supramodal regions in frontal and parietal lobes that carry out higher-order cognition, such as executive function, on the other (1-3). However, to fully understand how the interactions of these widespread brain systems give rise to conscious information processing, it is necessary to factor out brain processes that are not intrinsic to consciousness (4). To this end, functional neuroimaging of individuals rendered unconscious unde deep anesthesia or after severe brain injury provides a unique window for demarcating unconscious processes, and conversely, shedding light on brain mechanisms that are essential for conscious information processing and cognition in the healthy brain.

In a series of studies, we asked whether a common mechanism underlies loss of information processing in unconscious states across different conditions, which could shed light on the brain mechanisms of conscious cognition. To address this question we brought together, for the first time, data from the same paradigm—a highly engaging auditory-only narrative—in three independent domains: anesthesia-induced unconsciousness, unconsciousness after brain injury, and individual differences in intellectual abilities during conscious cognition.

Despite a growing number of anesthesia studies, it remains unknown how loss of consciousness affects synthesis of information across sensory and higher-order brain systems. To date, the majority of functional Magnetic Resonance Imaging (fMRI) studies of anesthesia have investigated the brain during a task- and stimulus-free condition, known as the “resting” state, because behavioral responses and eye opening are impaired by sedation prior to loss of consciousness (5), which render traditional experimental paradigms that probe complex information processing impossible to implement. However, because resting state studies do not use sensory stimulation, they cannot shed light on how the synthesis of external information breaks down from loss of consciousness. Several studies have used simple psychophysical stimuli and, therefore, have limited their investigation to well-circumscribed responses in sensory-specific cortex (6). In the auditory domain, studies have used simple auditory stimuli to investigate the limits of auditory processing during anesthetic-induced sedation. Following light anesthesia with sevoflurane, activation to auditory word stimuli relative to silence was preserved in bilateral superior temporal gyri, right thalamus, bilateral parietal, left frontal, and right occipital cortices (7). Parallel results have been found with both propofol and the short-acting barbiturate thiopental, suggesting that basic auditory processing remains intact during reduced or absent conscious awareness (6, 8-10). By contrast, light anesthesia impairs more complex auditory processing (11-12). For example, one study (13) showed that the characteristic bilateral temporal-lobe responses to auditorily presented sentences were preserved during propofol-induced sedation, whereas ‘comprehension-related’ activity in inferior frontal and posterior temporal regions to ambiguous versus non-ambiguous sentences was abolished. However, this study did not achieve the unconscious state due to low anesthetic doses. Thus, to date, no anesthetic study has directly investigated how the loss of consciousness affects the processing of a complex, real-world narrative across sensory-driven and higher-order brain systems.

Another group of individuals—patients who lose consciousness after severe brain injury—stand to shed light on the brain mechanism affected by loss of consciousness. Following serious brain injury, a proportion of patients manifest disorders of consciousness (DoC) and exhibit very limited responsivity to commands administered at the bedside by the clinical staff. If entirely behaviorally non-responsive, they are thought to lack consciousness—be in a vegetative state (VS) (14)—or, if they have reproducible but inconsistent willful responses, to be in a minimal conscious state (MCS) (15). The clinical, behavioral assessment of behaviorally non-responsive patients is particularly difficult and can result in high misdiagnosis rate (41%) (16). Studies show that, despite the apparent absence of external signs of consciousness, a significant minority of patients (∼19%) (17-19), thought to be in a VS, can demonstrate conscious awareness by willful modulation of their brain activity (20-26), a phenomenon captured by the recently proposed term ‘cognitive motor dissociation’ (CMD) (27). In the present study, to circumvent the limitations of behavioral testing and ensure that patients categorized as unconscious showed no willful brain responses, each patient underwent an fMRI-based assessment with a previously established command-following protocol for detecting covert awareness (22, 28). Similarly to the deep anesthesia context, experimental paradigms that probe the processing of complex external information have, until recently, not been implemented in DoC patients (29-32). Although, the disrupted brain mechanism in patients who are genuinely unconscious has been studied in the resting state paradigm (33-39), this, by default, cannot help to elucidate fully the mechanisms underlying loss of information processing in severely brain-injured unconscious patients.

The inherent limitations in testing unconscious individuals and the absence of identical sensory stimulation paradigms in anesthesia and severe brain injury investigations has hindered understanding of common mechanisms underlying loss of information processing across these conditions. To address this knowledge gap, in two different studies, we used the same paradigm and a novel approach (30) for measuring complex information processing in unconsciousness from deep anesthesia and severe brain injury, as participants freely listened to richly evocative stimulation in the form of a plot-driven narrative—a brief (5 minute) auditory-only excerpt from the kidnapping scene in the movie ‘Taken’. This approach circumvents traditional limitations by requiring neither behavioral response nor eye opening, and, importantly, elicits both sensory and fronto-parietal brain responses that are known to support high-order cognition, such as executive function (40-47). By their very nature, engaging narratives are designed to give listeners a common conscious experience driven, in part, by the recruitment of similar executive processes, as each listener continuously integrates their observations, analyses and predictions, while filtering out any distractions, leading to an ongoing involvement in the story’s plot. We have previously shown (30-32) that when different individuals freely listen to the same narrative, stereotyped changes of brain activity across these frontal and parietal cortical regions are observed, which reflect a robust and similar recruitment of executive function across different individuals. Thus, this paradigm is particularly suited for investigating the extent of information processing in behaviorally non-responsive individuals in unconscious states.

Conversely, we asked whether the principles of information processing revealed by the anesthesia and severe brain-injury studies could predict conscious cognitive performance, an independent domain that relies on continuously efficient processing of external information. Understanding individual differences in intellectual abilities is profoundly important as it may, in the future, help facilitate their enhancement, yet the underlying brain mechanisms remain poorly understood. Previous studies have suggested that functional connectivity within the fronto-parietal network during executive or cognitive tasks is related to individual differences in intelligence (40). This approach has been useful in identifying functionally segregated neural correlates of intelligence, i.e., the fronto-parietal network, but it does not reflect the role of sensory-driven networks or of their interactions with higher-order systems.

In the first study, we asked how information processing across the auditory and fronto-parietal systems during the story was affected by loss of consciousness in deep anesthesia in healthy participants (N=16). In the second study, we tested whether the insights gleaned from the anesthesia study could generalize to loss of consciousness after severe brain injury, in a group of patients (N=11) with disorders of consciousness that underwent fMRI scanning during the same audio story as healthy participants from study one. In the third study, we investigated how the cognitive performance of the individuals from the anesthesia study (N=14) independently-measured with a cognitive battery weeks after the sedation study related to their synthesis of complex sensory information between auditory and fronto-parietal systems during the audio-story task.

## Results

### Information processing under deep anesthesia

To measure information processing during the story, we adopted a previously established method using the same audio story (30), where we showed that the extent of stimulus-driven cross-subject correlation provided a measure of regional stimulus-driven information processing (Figure 1A-C). In the wakeful condition of the anesthesia study, we observed widespread and significant (p<0.05; FWE cor) cross-subject correlation between healthy participants within sensory-driven (primary and association) auditory cortex, as well as higher-order frontal and parietal regions (Figure 1D), consistent with Naci et al. (2017) (30). By contrast, in deep anesthesia, the significant (p<0.05; FWE cor) cross-subject correlation was limited to the auditory cortex, with the exception of two small clusters in left prefrontal and right parietal cortex (Figure 1E), suggesting that the processing of sensory information was preserved in the sensory, but almost entirely abolished in fronto-parietal regions.

**Figure 1.**
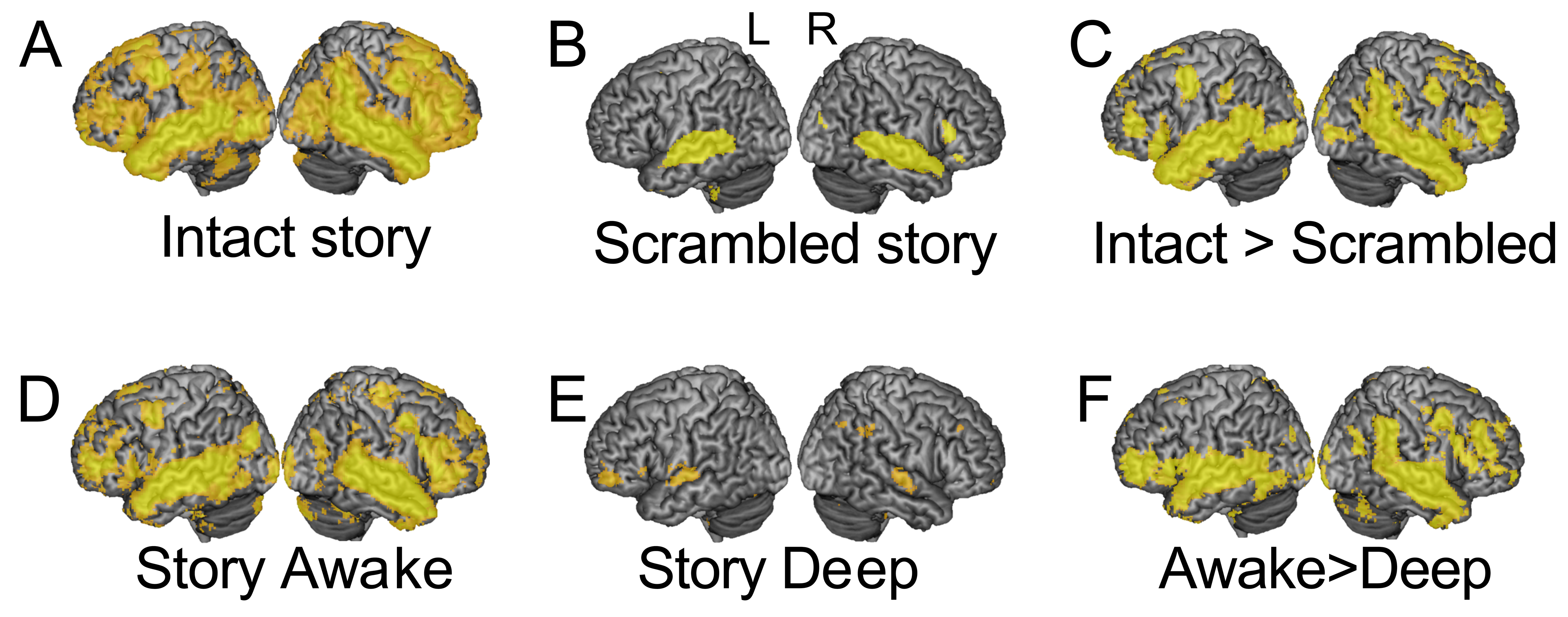
Brain-wide inter-subject correlation of neural activity during the audio story. (A) The audio story elicited significant (p<0.05; FWE cor) inter-subject correlation across the brain, including frontal and parietal cortex, thought to support executive function. (B) The baseline elicited significant (p<0.05; FWE cor) inter-subject correlation within primary and association auditory cortex. A small cluster was also observed in right inferior prefrontal cortex. None was observed in dorsal prefrontal and parietal cortex. (C) The audio story elicited significantly (p<0.05; FWE cor) more inter-subject correlation than the auditory baseline derived from the same stimulus, in parietal, temporal, motor, and dorsal/ventral frontal/prefrontal cortex. A, B, C, adapted with permission from Naci et al. (2017) (**30**). (D) The audio story elicited significant (p<0.05; FWE cor) inter-subject correlation across the brain, including frontal and parietal cortex, in the wakeful state of the anesthesia study. (E) In the deep anesthesia state, significant (p<0.05; FWE cor) inter-subject correlation was limited to the auditory cortex with the exception of two small clusters, one in left prefrontal and the other in right parietal cortex. (F) The intact audio story elicited significantly (p<0.05; FWE cor) more cross-subject correlation than the scrambled story bilaterally in temporal, ventral prefrontal and frontal cortex, and further in parietal, motor, and dorsal frontal and prefrontal cortex in the right hemisphere. Warmer colors depict higher t-values of cross-subject correlation. Warmer colors depict higher t-values of inter-subject correlation.

Subsequently, we investigated the impaired brain mechanism underlying loss of information processing in these higher-order regions. Current theories of consciousness (48-50), such as the Integrated Information Theory (IIT), propose that conscious cognition relies on the brain’s capacity to efficiently integrate information across different specialized systems (48), suggesting that both interconnectedness and functional differentiation of brain systems are important for information processing. However, different putative mechanisms are consistent our results, including reduced/abolished connectivity among distinct brain systems (49) and loss of functional differentiation (48) (i.e., homogeneous connectivity across them). To directly investigate the underlying mechanism, we distinguished four possible impairment patterns consonant with theories of consciousness that could explain impaired information processing in deep anesthesia: 1) a loss of long-range connectivity between auditory and fronto-parietal networks (Figure 2A); 2) a loss of connectivity between areas within each network, e.g., between frontal and parietal regions (Figure 2B); (3) a combination of 1 and 2 (Figure 2C); and, (4) a loss of differentiation between auditory and fronto-parietal networks (Figure 2D).

**Figure 2.**
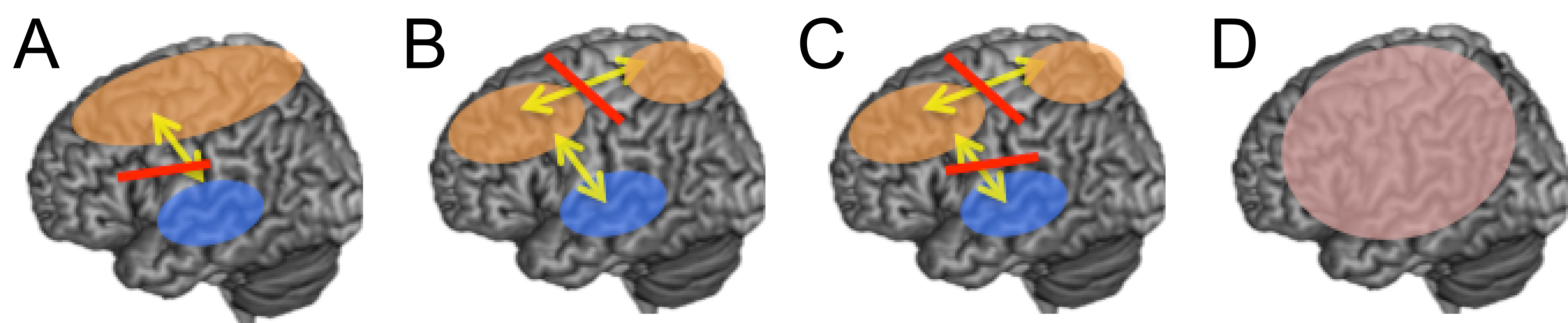
Candidate patterns of connectivity perturbations caused by deep propofol anesthesia. (A) Loss of long-range connectivity between different networks; (B) Loss of long-range connectivity within a specific network, e.g., between frontal and parietal regions; (C) A combination of patterns in A and B; (D) Loss of functional differentiation between different brain networks.

### The global effect of anesthesia on brain networks’ connectivity

Initially, we investigated how deep propofol anesthesia perturbed the patterns of global connectivity. During the audio story, a two-way ANOVA with factors connectivity type (within, between) and state (wakeful, deep anesthesia) showed that connectivity across networks increased significantly (main effect of state: F(16)=8.57; p=0.01) (Figure 3A-B) in deep anesthesia relative to wakefulness. Connectivity *between* increased more than *within* networks (interaction effect, state x connectivity type: F(15)=5.58; p<0.05) (Figure 3E), driven by a significant increase in the *between* network connectivity (t(15)=3.82, p<0.005) and no overall change in the *within* connectivity (see SI for complete results; Figure S3, S4). By contrast, during the resting state, deep anesthesia showed the opposite effect on between and within network connectivity, with a larger impact on the *within* relative to *between* network connectivity (interaction effect, state x connectivity type: F(15)=5.4; p<0.05) (Figure 3C-D). Connectivity *within* was significantly reduced, but no changes were observed in deep anesthesia in the *between* network connectivity (see SI for complete results).

**Figure 3.**
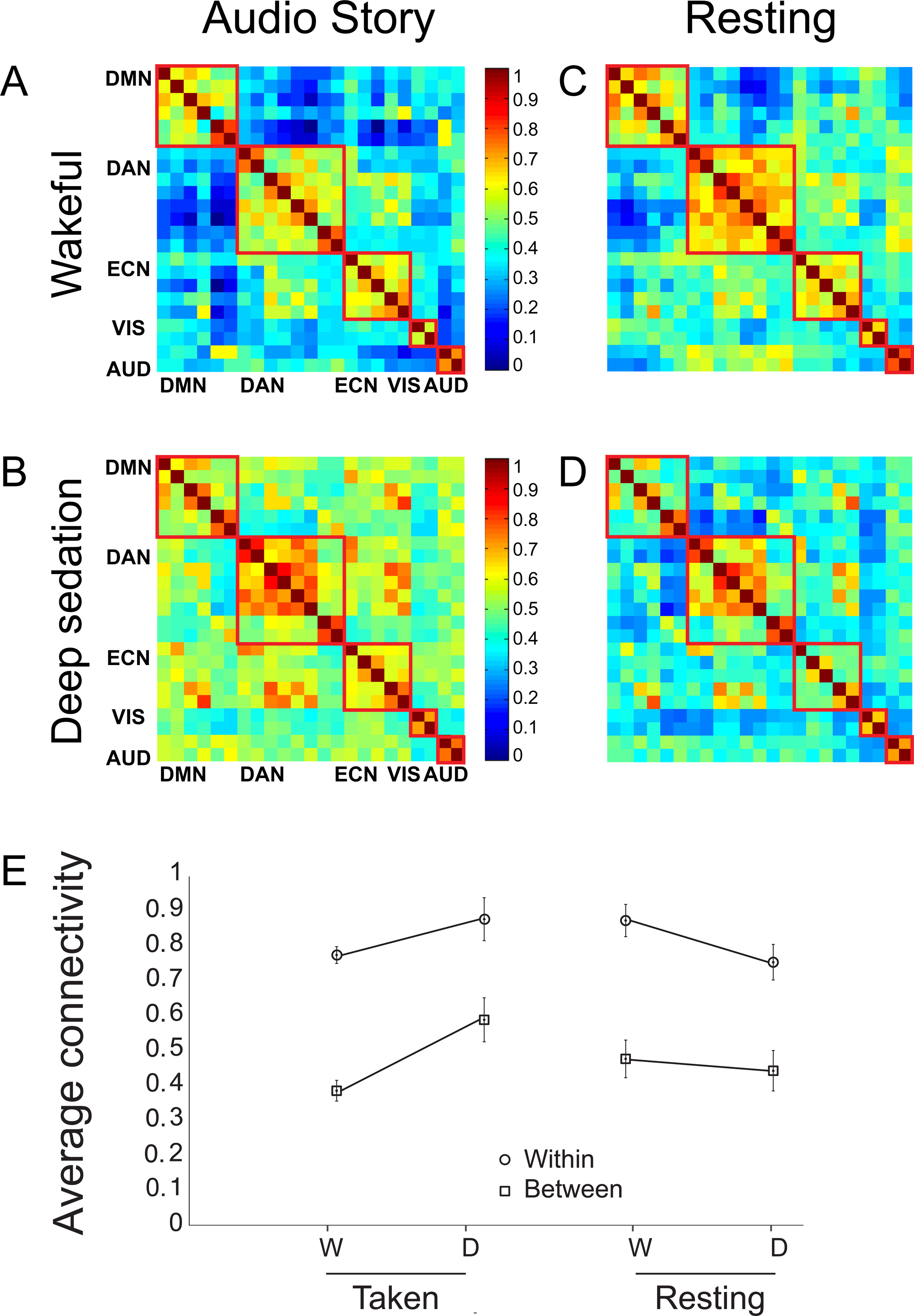
Global *within*- and *between*-network functional connectivity perturbations by deep propofol anesthesia. (A–D) Functional connectivity matrices for five brain networks in the story and resting state conditions, in the wakeful and deep anesthesia states. Each cell represents the correlation of the time-course of one region of interest (ROI) with another, or itself (in the center diagonal). Cells representing correlations of ROIs within each network are delineated by red squares. Warm/cool colors represent high/low correlations, as shown in heat-bar scale. (E) Average connectivity (z values) *within-* and *between-* networks in the wakeful (W) and the deep anesthesia (D) states, during the story and resting state conditions. DMN/DAN/ECN/VIS/AUD=Default Mode/Dorsal Attention/Executive Control/Visual/Auditory network.

A direct comparison between the audio story and resting state confirmed that anesthesia affected connectivity in the two conditions in opposite directions. A two-way ANOVA with factors condition (audio story, resting state) and state (wakeful, deep anesthesia) showed a condition x state interaction [F(15)=7.01; p<0.05] that was driven by an overall connectivity reduction during the resting state and connectivity increase during the audio story in deep anesthesia. The effects were the same when functional differentiation was measured as the ratio of *between-* to *within-*network connectivity.

These suggest that, when the brain is at rest, reduced connectivity within brain networks rather than loss of functional differentiation between them, characterizes the unconscious state. By contrast, when the brain is exposed to complex naturalistic stimuli from the environment, reduced functional differentiation between brain networks leads to loss of information processing in the unconscious state. However, these results must be interpreted with caution, in light of the consistent block order in deep sedation.

### The effect of anesthesia on auditory and fronto-parietal networks’ connectivity

Next, we asked specifically whether reduced functional differentiation between the auditory and fronto-parietal networks drove the loss of information processing in the fronto-parietal regions during the story. Consistent with effects at the whole-brain level, we found a significant increase in the AUD–DAN and AUD–ECN connectivity [t(15)=2.6, p<0.05; t(15)= 4.98, p<0.0005, respectively] (Figure 4A-D), or a significant reduction of the functional differentiation between the AUD and DAN, ECN in deep anesthesia relative to wakefulness. By contrast, in the resting state, connectivity between these networks was not affected by sedation (Figure 3). These results suggested that reduced functional differentiation between the auditory and fronto-parietal networks leads to loss of external information processing in the unconscious state. Conversely, they suggested that the functional differentiation between the auditory and fronto-parietal networks underlies conscious processing of complex auditory information.

To test specifically whether functional differentiation in the conscious state would be driven by the complex features of the audio story, including it’s narrative, rather than merely the presence of external stimulation, we compared the AUD–DAN and AUD– ECN pairwise connectivity in wakeful individuals during the audio story with those in the two baseline conditions, a scrambled version of the story that retained the sensory features but was devoid of the narrative, and the resting state. During the intact audio story, functional connectivity between the AUD and DAN, ECN was significantly lower than in the scrambled story [AUD–DAN: t(14)=-11.2, p<0.0001; AUD–ECN: t(14)=- 10.62, p<0.0001], and than in the resting state [AUD–DAN: t(14)=-7.3, p<0.0001; AUD– ECN: t(14)=-2.7, p<0.05] (Figure 5A-H). These results suggested that the processing of the high-order features of story, including its narrative, drove functional differentiation between the auditory and fronto-parietal networks in wakeful individuals.

**Figure 4.**
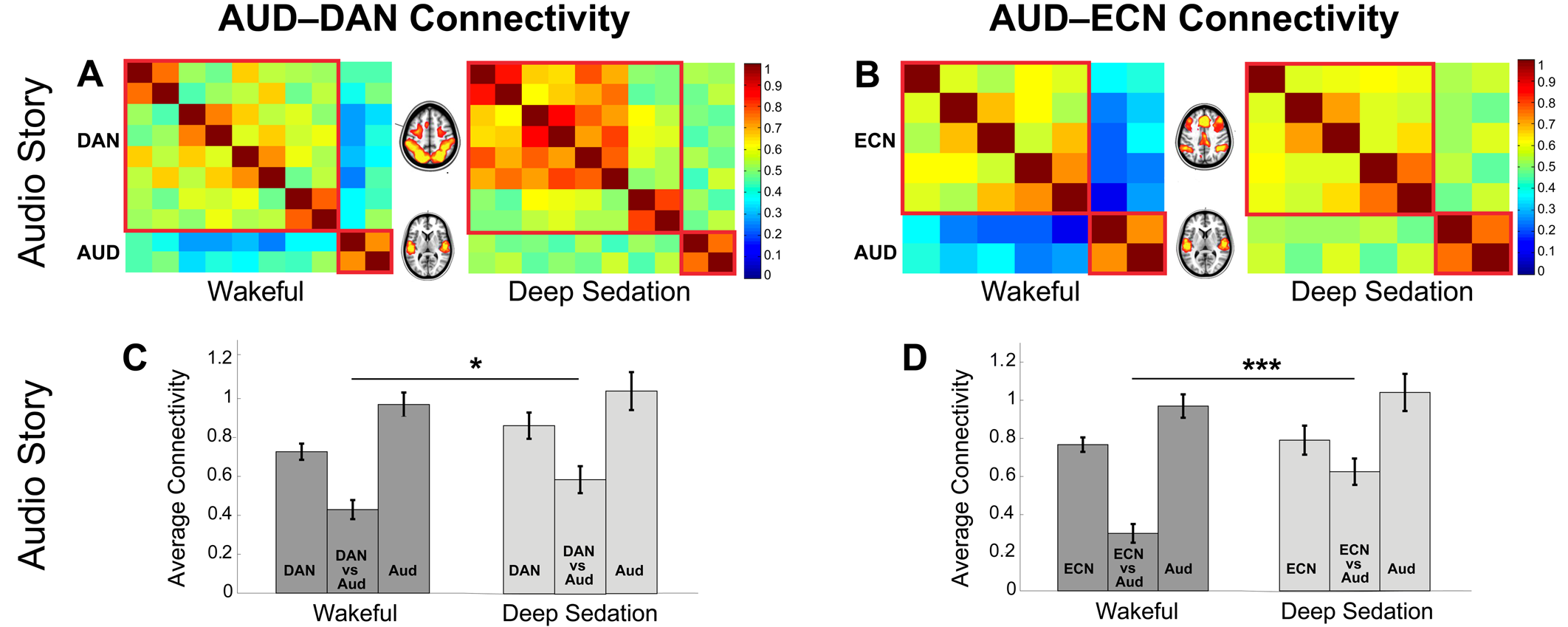
Perturbations of auditory and fronto-parietal connectivity by deep propofol anesthesia in the audio story condition. Only connectivity *between* the AUD and DAN/ECN, respectively, was significantly modulated by propofol, showing a significant reduction of functional differentiation between sensory and higher-order networks in deep anesthesia relative to wakefulness. (A–B) Functional connectivity matrices for ROIs comprising the DAN and AUD (A)/ECN and AUD (B) networks in the wakeful and deep anesthesia states of the audio story condition. (C–D) Average connectivity (z-values) *within* and *between* the DAN and AUD (C)/ECN and AUD (D), in the wakeful and deep anesthesia states.

**Figure 5.**
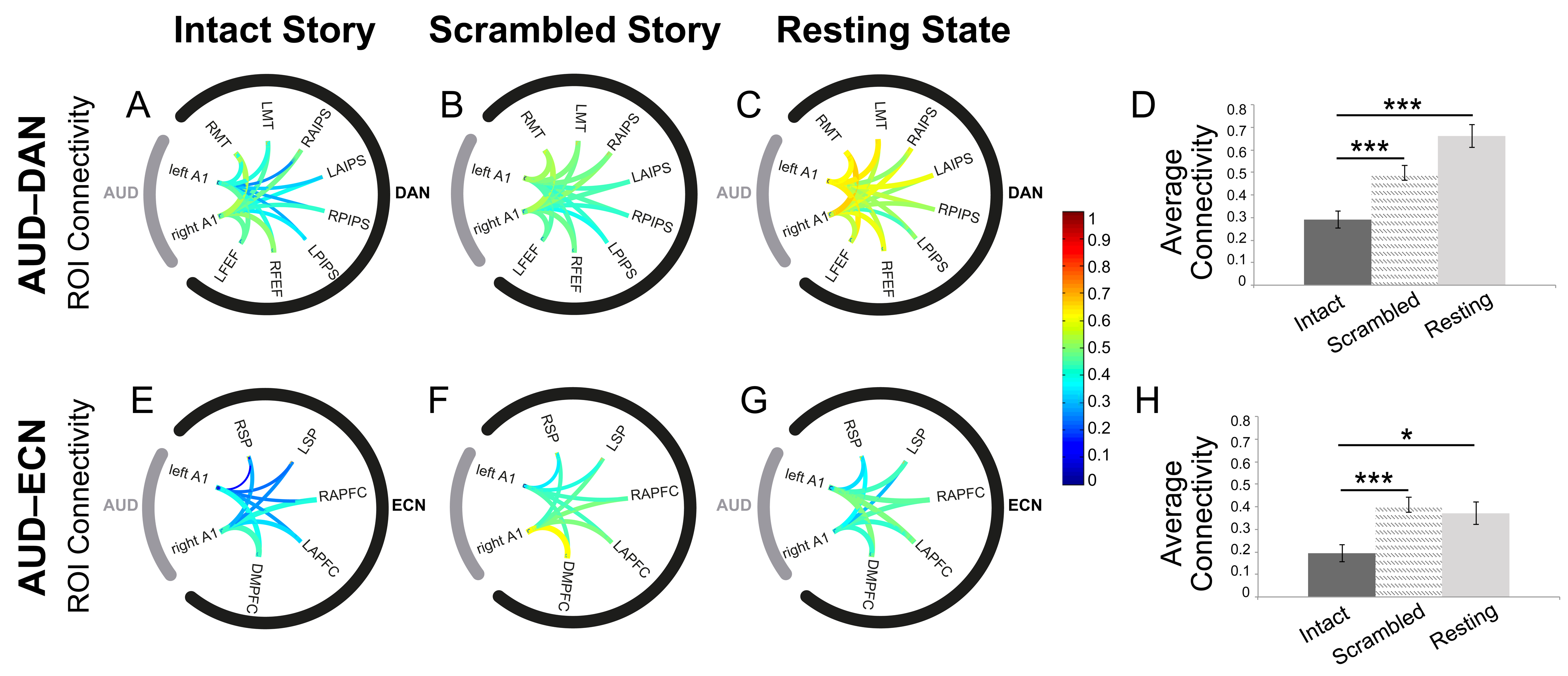
Functional connectivity between the auditory and fronto-parietal networks in healthy wakeful individuals, during the audio story and baseline conditions. Connectivity between the auditory and fronto-parietal networks was significantly modulated by the presence of complex meaningful stimuli, with the functional differentiation between the AUD and DAN/ECN increasing significantly in the audio story as compared to the scrambled story and resting state baseline conditions. (A–C) Connectivity between the ROIs within the AUD and DAN networks in the intact story (A), scrambled story (B), and resting state (C) baseline. (D) Average AUD–DAN connectivity (z values) for each condition. (E–G) Connectivity between the ROIs within the AUD and ECN networks in the intact story (E), scrambled story (F), and resting state (G) baseline. (H) Average AUD–ECN connectivity (z values) for each condition. A1: Primary auditory cortex; LFEF: Left frontal eye field; RFEF: Right frontal eye field; LPIPS: Left posterior IPS; RPIPS: Right posterior IPS; LAIPS Left anterior IPS; RAIPS: Right anterior IPS LMT: Left middle temporal area; RMT: Right middle temporal area; DMPFC: Dorsal medial PFC; LAPFC: Left anterior prefrontal cortex; RAPFC: Right anterior prefrontal cortex LSP: Left superior parietal; RSP: Right superior parietal.

### The effect of severe brain injury on the auditory and fronto-parietal networks’ connectivity

Results from both conscious and unconscious conditions in the previous study suggested that functional differentiation between the auditory and fronto-parietal networks underlined conscious processing of complex auditory information. In the next study, we further tested this claim in severe brain-injury, which served as an independent manipulation of consciousness.

The structural profiles and full behavioral description of the convenience sample of brain-injured patients (N=11) are shown in Figure S1, Table S2, S3. Patients who showed willful brain responses in the independent command-following assessment (22, 28) were considered covertly aware and labeled DoC+ (N=6), and those who showed no signs of conscious awareness were labeled as DoC-(N=5) (Figure 6), for subsequent analyses. Similarly to conscious individuals (Figure 5), we expected DoC+ patient group to show a heightened differentiation/down-regulation of the AUD and DAN, ECN pairwise connectivity during the audio story relative to resting state baseline connectivity. By contrast, we did not expect a down-regulation of the connectivity between these networks during the audio story in DoC-patient group.

**Figure 6.**
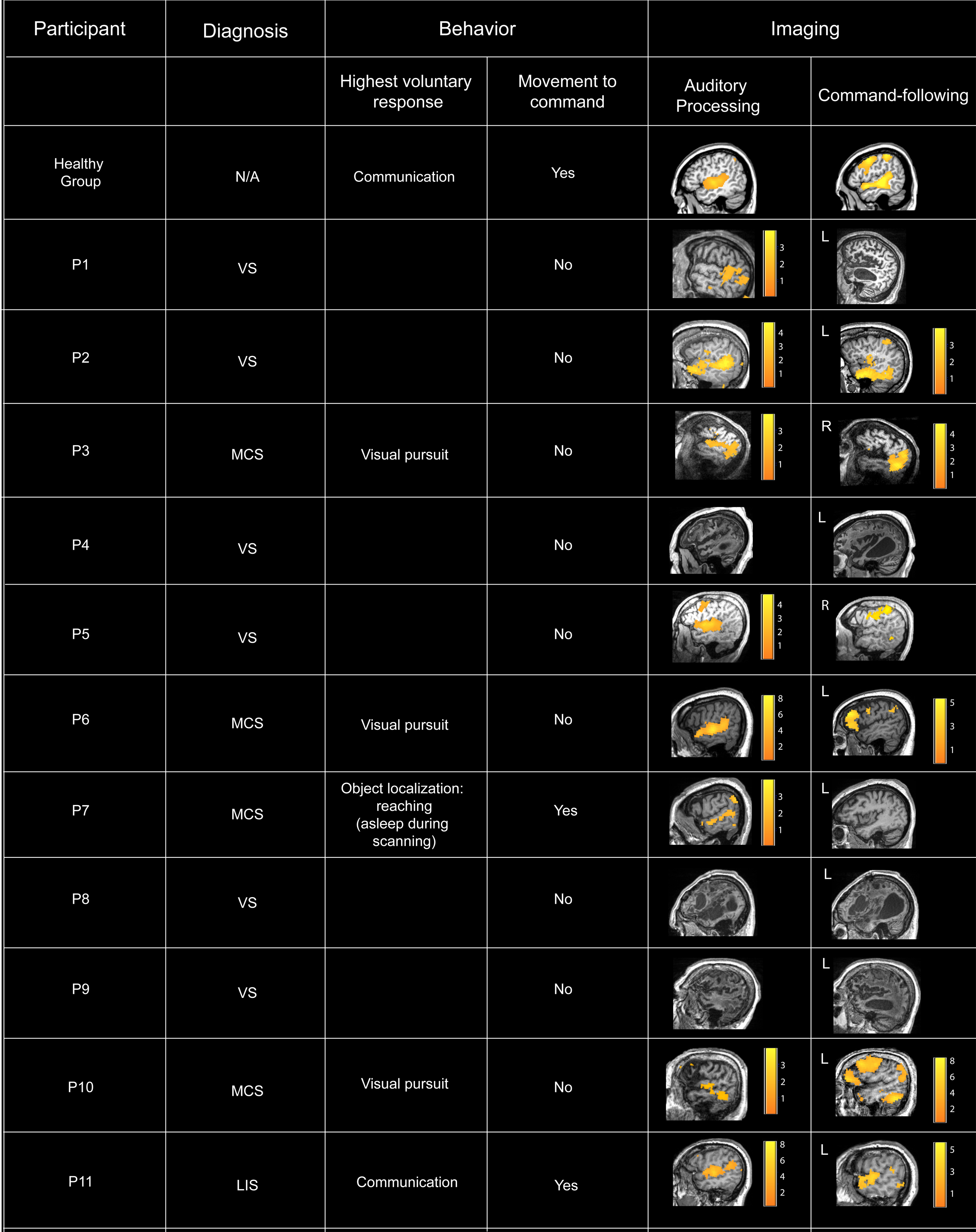
Summary of DoC patients’ clinical and fMRI assessment data. *Auditory processing.* In the fMRI assessment, three patients clinically diagnosed to be in a VS did not show evidence of auditory processing. The other eight patients who showed evidence of auditory processing, two patients clinically diagnosed as VS did not show evidence of brain-based command-following, and the other six, including two diagnosed as VS, showed evidence of brain-based command-following, and thus, of covert awareness. *Command-following*. 6/11 patients followed task commands by willfully modulating their brain activity as requested, and thus, provided evidence of conscious awareness. Two of these (P2, P5) presented a CMD profile, or a behavioral diagnosis of VS that was inconsistent with their positive fMRI results. 5/11 patients showed no evidence of willful responses in the fMRI command-following task, and, thus, provided no neuroimaging evidence of awareness. One (P7) showed no neuroimaging evidence of awareness despite an MCS diagnosis, due to falling asleep in the scanner for the entirely of the session (Materials and Methods).

The DoC+ group showed a significant down-regulation of the auditory and fronto-parietal networks connectivity in the audio story relative to the resting state [AUD–DAN: t(5)=-1.9, p=0.05; AUD–ECN: t(5)=-3.6, p<0.5] (Figure 7A-H, E-L), and was significantly different from the DoC- (N=5) group, who did not show this effect (Figure 7C-J, E-L) [AUD–DAN: t(8)=-3.6, p<0.01; AUD–ECN: t(9)=-3.4, p<0.01]. The predicted effect pattern was also observed for individual patients, with 5/6 DoC+ patients showing a down-regulation of the connectivity between AUD and DAN, ECN (Figure 7F, L). The effect in the DoC+ group was consistent with the effect observed in healthy conscious individuals (Figure 5). By contrast, the DoC-group showed significantly enhanced AUD–DAN connectivity during the audio story relative to resting state [t(4)=4.48; p<0.05] (Figure 7E). This was consistent with the up-regulation of the AUD– DAN connectivity observed in the anesthesia-induced unconscious state in the previous study.

**Figure 7.**
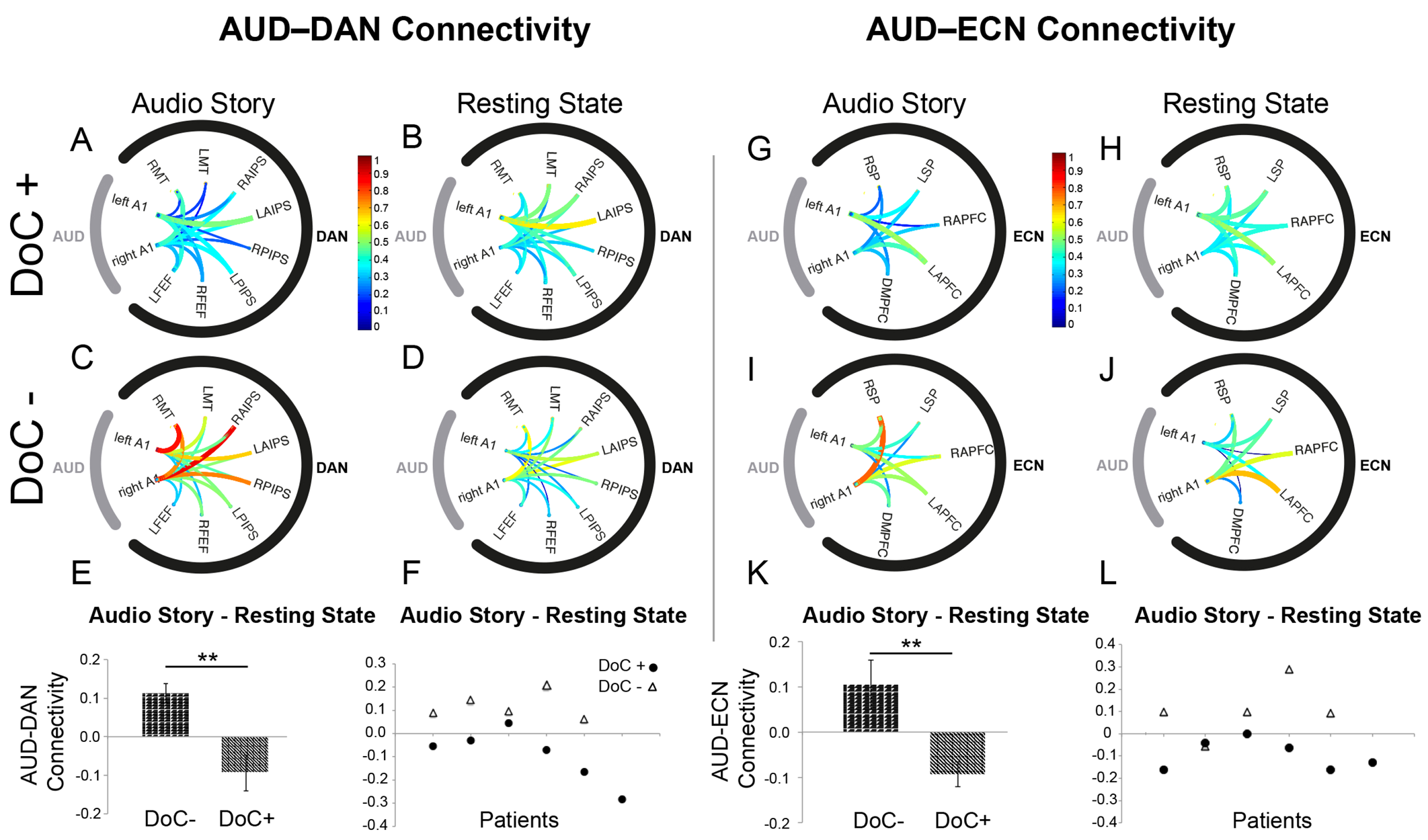
Modulation of auditory to fronto-parietal connectivity by meaningful stimulation in DoC patients. Similarly to healthy individuals, connectivity between the auditory and fronto-parietal networks in DoC+ patients was significantly modulated by the presence of complex meaningful stimuli, with the functional differentiation between the AUD and DAN/ECN increasing significantly in the audio story as compared to the resting state baseline condition. (A–D) Connectivity between the ROIs within the AUD and DAN networks, during the audio story and resting state baseline, in the DoC+ (A–B) and DoC- (C–D) patients. (E) Differential averaged AUD–DAN connectivity (z values) for each patient group. (F) Differential averaged AUD–DAN connectivity (z values) for each individual patient. (G–J) Connectivity between the ROIs within the AUD and ECN networks, during the audio story and resting state baseline, in the DoC+ (G–H) and DoC- (I–J) patients. (K) Differential averaged AUD–ECN connectivity (z values) for each patient group. (L) Differential averaged AUD–ECN connectivity (z values) for each individual patient. A1: Primary auditory cortex; LFEF: Left frontal eye field; RFEF: Right frontal eye field; LPIPS: Left posterior IPS; RPIPS: Right posterior IPS; LAIPS Left anterior IPS; RAIPS: Right anterior IPS LMT: Left middle temporal area; RMT: Right middle temporal area; DMPFC: Dorsal medial PFC; LAPFC: Left anterior prefrontal cortex; RAPFC: Right anterior prefrontal cortex LSP: Left superior parietal; RSP: Right superior parietal.

### Network connectivity and individual differences in conscious cognition

Taken together, the results from the two previous two studies suggested that heightened differentiation between the auditory and fronto-parietal networks supports conscious processing of complex auditory information, and more broadly, conscious cognition. In the third study, we further tested this claim directly, by asking whether it predicted individual differences in cognitive performance. We assessed the cognitive performance of a participant subset (14/16) from the anesthesia study, who came back to the laboratory weeks later, with a battery comprising 12 cognitive tests (51) that measured short-term memory, reasoning, and verbal acuity (SI, Table S4). Based on converging results from studies 1 and 2, we expected stronger differentiation between the auditory and fronto-parietal networks during the audio story to predict stronger cognitive performance, or, a negative relationship between the AUD and DAN, ECN connectivity and the independently measured cognitive performance in the same individuals.

The individuals’ AUD–DAN connectivity during the audio story was significantly negatively correlated (r = -0.66; p<0.05) with their cognitive performance in the verbal acuity (Figure 8 A-B) component of the battery, which accounted for the variance of tasks that used verbal stimuli (i.e., digit span, verbal reasoning, color-word remapping; Supporting Information). The AUD–DAN connectivity did not predict performance in the other two components, and the AUD–ECN connectivity did not predict performance in any of the three (Figure 8A, C). Pairwise connectivity between these networks in the resting state did not predict cognitive performance in any of the domains (Figure 8A). Further, we found no relationship between the connectivity of the AUD and default mode network (DMN), included as a control high-order network, and cognitive performance.

**Figure 8.**
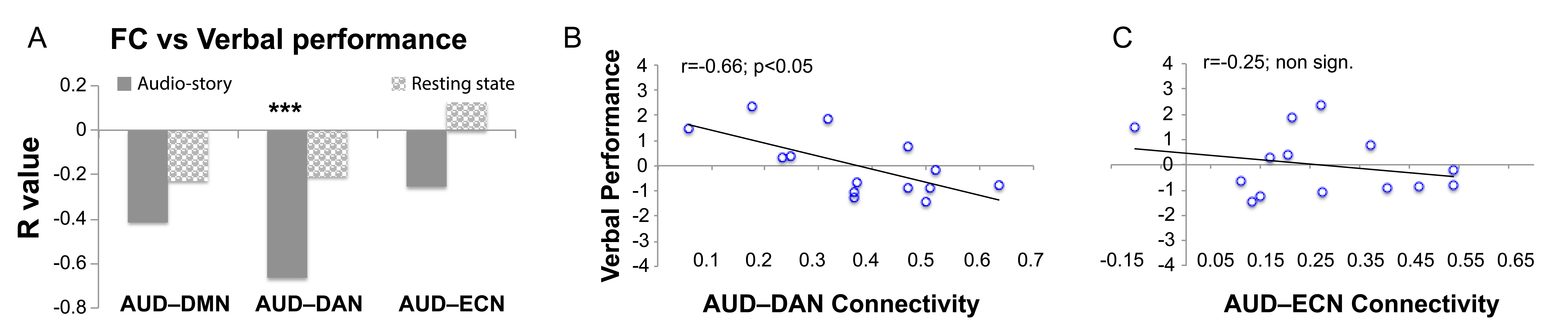
The relationship between brain network connectivity during the audio story and independently-measured cognitive performance. The functional connectivity between AUD and DAN, but not ECN (or DMN, used here as a high-level control network) during the story, and not the resting state baseline condition, was significantly inversely related to verbal performance. (A) Group-averaged correlation between the functional connectivity (FC) of the AUD and the DMN/DAN/ECN networks during the audio story and resting state conditions and verbal performance. (B–C) For each participant, the correlation between their AUD–DAN (B)/AUD–ECN (C) connectivity during the story and their verbal performance is displayed.

In summary, the results of the third study converged with the other two, and suggested that the extent to which the functional responses of the auditory and fronto-parietal networks to complex auditory stimuli dissociated from one another predicted independent cognitive performance in the verbal domain, and thus, may be a determining factor in individual differences in verbal acuity.

Figure 8. The relationship between network connectivity during the audio story and independently-measured cognitive performance.

## Discussion

In a series of studies, we asked whether a common mechanism underlies loss of information processing in unconscious states across different conditions, which could shed light on the brain mechanisms of conscious cognition. To this end, for the first time, we brought together two very disparate conditions where consciousness is lost—deep anesthesia and severe brain injury—to investigate the modulation of functional connectivity between the auditory and fronto-parietal networks by identical complex stimulation in identical paradigms. Subsequently, we tested whether findings from these studies predicted individual differences in intellectual abilities during conscious information processing.

### Common mechanism for loss of information processing in unconsciousness during anesthesia and after severe brain injury

We use a novel approach (30-32), to measure external information processing in response to richly evocative stimulation portraying real-world events, during deep anesthesia and severe brain injury. In the anesthesia study, we found that the processing of the story information was preserved in auditory cortex, but almost entirely abolished in fronto-parietal regions. Deep anesthesia led to a significant reduction in the functional differentiation of several networks across the brain, and specifically, between the auditory and fronto-parietal networks, during the story condition. These results suggested that anesthesia impaired the processing of complex external information in fronto-parietal regions by eroding their functional differentiation from sensory (e.g., auditory) systems, and not by impairing connections between or within them. Propofol was used here as a common anesthetic agent, and future studies that employ the same paradigm across different agents will help elucidate whether specific agents vary in their effect on connectivity during complex stimulation. Our results are consistent with previous findings from resting state studies, suggesting that anesthesia reduces the repertoire of discriminable brain states (52-53), and that during loss of consciousness global synchrony impairs information processing by leading to a breakdown of causal interactions between brain areas (54-56). Further, they are consistent with resting state studies using sleep-induced altered states of consciousness, which show that hyper-synchrony perturbs the feed-forward propagation of auditory information (57), as well as feedback projections (58), and more broadly, the stable patterns of causal interactions in response to external stimulation across the brain (59). While these previous resting state studies suggest that global synchrony breaks down causal interactions, the investigation of causal cortico-cortical interactions was outside the scope of this work. We did not find an effect of deep sedation on thalamo-cortical connectivity in any of the five brain networks (SI), and while outside of our scope here, a potential causal role of thalamic inputs to cortico-cortical connectivity in deep sedation remains to be investigated further.

Our findings from the resting state condition in deep anesthesia manipulation agree with a previously reported reduction of brain connectivity in deep propofol anesthesia during the resting state (49, 54, 60), in particular with a reduction of connectivity *within* the default-mode 60-62; but, see 53, 63), and the executive control networks (60, 64). Although consistency with previous studies suggests otherwise, we note that differences between the story and resting state conditions in deep sedation must be interpreted with caution, in light of the consistent block order. Nevertheless, the results from the sensory stimulation condition reveal a different mechanism underlying the loss of external information processing than suggested by resting state studies. First, in the audio-story condition, the connectivity within networks was affected by sedation in the opposite direction to the resting state. Second, in the resting state condition, we observed no effect of deep anesthesia on connectivity *between* distinct networks, which, by contrast, increased significantly during the auditory stimulation condition suggesting loss of functional differentiation across the cortex. Another type of stimulation—transcranial magnetic stimulation 3TMS)—has previously been used to directly perturb the cortex in unconscious states and demonstrate that responses across the cortex become undifferentiated from one another (55, 65). In summary, these results suggest that deep anesthesia affects the brain differently when it is exposed to complex external stimulation relative to rest, with the stimulus-evoked feed-forward processing cascade being echoed undifferentiated throughout the brain, thus overcoming the inhibitory effect of propofol on neural connectivity that has been reported in resting state studies (66).

Similarly to deeply anesthetized unconscious individuals, severely brain-injured patients who were not consciously aware during the study showed significantly reduced differentiation between the auditory and fronto-parietal networks during the story relative to their resting baseline. Conversely, similarly to healthy wakeful individuals, severely brain-injured patients who were covertly aware showed the opposite effect: significantly enhanced differentiation between the auditory and fronto-parietal networks during the story relative to their resting baseline. The modulation of the sensory to higher-order networks’ relationship by meaningful environmental stimuli in severely brain-damaged (albeit conscious), patients suggests this is a fundamental feature of conscious processing that is resilient to substantial metabolic dysfunction following brain injury (67). We caution that our results do not suggest that each DoC+ patient understood the story similarly as healthy individuals. Brain-injured patients fluctuate vastly in arousal and, thus, even if individual DoC+ patients discussed here retained functional brain architecture to support covert conscious awareness, the absence of a sensory baseline and individual-level statistics, render it impossible to ascertain the extent of understanding in individual patients.

Previous studies that have compared anesthetized and unconscious brain-injured patients have highlighted that, similarly to the effect of common anesthetic agents including propofol (60, 68), brain dysfunction in this population is prevalent within the fronto-parietal network (69-70). They have indicated preserved sensory processing (e.g., responses to noxious stimulation, auditory or speech perception) in the absence of higher-order components (e.g., neural evidence of pain perception, language comprehension) (5), and suggested that disconnection between sensory and fronto-parietal systems is common to both populations. By contrast, our findings suggest that, when the brain is exposed to complex external information, these systems do not disconnect from one another in these unconscious states, as previously suggested by aforementioned resting state studies. Rather, our findings demonstrate that the erosion of functional differentiation among these systems underlies impaired information processing when consciousness is lost.

Although loss of consciousness is common to both deeply anesthetized and some severe brain-injured patients, these two populations differ greatly. In the former no structural changes occur, and the functional brain response is altered pharmacologically. In the latter, an array of structural damage, greatly varying across patients, is present and affects altered brain responses, leading to complete functional loss in some domains and potential functional re-organization and preservation in others. Given the large differences between these two, the similarity of the functional response to previously validated targeted stimulation (30) across these populations provides strong evidence for a common mechanisms underlying loss of information processing in the unconscious state. These results are consistent with current theories of consciousness, which suggest that it requires both differentiation and integration of information in neural circuits (48, 54, 71), and elucidate the underlying brain mechanism by showing the critical role of functional differentiation between sensory and higher-order systems when information processing is required.

### Mechanism for conscious information processing and cognition

The third study further confirmed the role of the functional differentiation between the sensory and higher-order systems in conscious cognition. Individuals who showed higher differentiation between the auditory and dorsal attention network (DAN) in response to the audio story had higher verbal acuity scores than individuals who showed lower differentiation. The story elicited a range of cognitive processes such as the orientation and modulation of attention to the saliency of incoming auditory inputs—a function subserved primarily by the DAN (72) — and language perception and comprehension, which corresponded to those engaged by the verbal acuity tasks of the cognitive battery. The functional relationship between the auditory and executive control network (ECN) was not predictive of cognitive performance, which is likely accounted for by the nature of the stimulus and fMRI paradigm which did not require behavioral response planning or monitoring—a function sub-served primarily by the ECN (73). There was no relationship between the auditory and fronto-parietal connectivity in response to the story and performance in the short-term memory or reasoning components of the cognitive battery, likely due to the story’s cognitive demands low loading on these components. We note that the verbal component of the cognitive battery, which comprised 12 tasks, accounted for the majority of variance in a subset of different tasks that used verbal stimuli (digit span, verbal reasoning, color-word remapping; full description in SI). Thus, in capturing a cross-section of processes employed in these different tasks, the verbal component represented a robust example of varied domain-specific processes, which are abstracted away from the demands of particular tasks. Therefore, although these results suggested that the relationship between brain connectivity and intelligence is domain-specific, future studies are required to further test the sensory–higher-order networks’ relationship and other cognitive domains/processes.

Further, these results agree with a previous proposal that the relationship between brain connectivity and intelligence is context specific (74). In contrast to the a-priory predicted relationship between these networks’ connectivity and intelligence during complex sensory stimulation, we found no relationship between them in the resting state. Notably, these results were predicted from two different populations where loss of from information in unconsciousness suggested a common mechanism for information processing during conscious cognition. Consistent with a recent emerging view in the field (75), they suggested that individual differences in intellectual abilities rely on the dynamic reconfigurations of connectivity in response to incoming sensory information, within a widespread system comprising sensory-specific and extra-modal cortices in fronto-parietal cortex.

In summary, findings herein suggest that the dissolution of functional differentiation is a common basis for loss of information processing across widely different conditions where consciousness is lost. Conversely, they suggest that the responsivity of sensory and higher-order brain systems to naturalistic external stimulation, especially through the diversification of their functional responses is an essential feature of conscious cognition and domain-specific intelligence.

## Material and Methods

### Participants

Ethical approval was obtained from the Health Sciences Research Ethics Board and Psychology Research Ethics Board of Western University. All experiments were performed in accordance with the relevant guidelines and regulations set out by the research ethics boards. All healthy volunteers were right-handed, native English speakers, and had no history of neurological disorders. The respective substitute decision makers gave informed written consent for each patient’s participation. They signed informed consent before participating and were remunerated for their time. 19 (18–40 years; 13 males) healthy volunteers, 11 (19-55 years; 5 males) DoC patients, and 14 (18–40 years;12 males) healthy volunteers participated in study 1, 2, and 3, respectively. Three volunteers (1 male) were excluded from data analyses of study 1, due to headphone malfunction or physiological impediments to reaching deep anesthesia in the scanner.

### Stimuli and Design

In study 1, a plot-driven audio story (5 minutes) was presented in the fMRI scanner to healthy volunteers and they were asked to simply listen with their eyes closed. A resting state scan (8 minutes) was also acquired, during which volunteers were asked to relax with their eyes closed and not fall asleep. A novel re-analysis of data from the scrambled story condition from Naci et al. (2017) (30) (SI) was performed, as a baseline condition with the intact audio story. In study 2, severely brain-injured patients were scanned as they listened to the same audio story as healthy volunteers, and also during the resting state. In study 3, 14/16 of volunteers from the anesthesia study completed a cognitive battery comprising 12 tasks based on classical cognitive psychology paradigms (www.CambridgeBrainSciences.com) (SI). The stimuli and design for each were reported in Hampshire et al. (2012) (51).

### Sedation procedure

fMRI data was acquired during the audio story and resting state conditions while participants were awake (non-sedated) and deeply anesthetized with propofol (Ramsay score 5) (76). Prior to acquiring fMRI data for the wakeful and deeply anesthetized states, 3 independent assessors (two anesthesiologists and one anesthesia nurse) evaluated each participant’s Ramsay level by communicating with them in person inside the fMRI scanner room, as follows. *Awake Non-sedated.* Volunteers were fully awake, alert and communicated appropriately. For the wakeful session, they were not scored on the Ramsay sedation scale, which is intended for patients in critical care settings or patients requiring sedation. During the wakeful audio story and resting state conditions, wakefulness was monitored with an infrared camera placed inside the scanner. *Deep anesthesia.* Intravenous propofol was administered with a Baxter AS 50 (Singapore). We used an effect-site/plasma steering algorithm in combination with the computer-controlled infusion pump to achieve step-wise increments in the sedative effect of propofol. The infusion pump was manually adjusted to achieve desired levels of sedation, guided by targeted concentrations of propofol, as predicted by the TIVA Trainer (the European Society for Intravenous Aneaesthesia, eurosiva.eu) pharmacokinetic simulation program. The pharmacokinetic model provided target-controlled infusion by adjusting infusion rates of propofol over time to achieve and maintain the target blood concentrations as specified by the Marsh 3 (77) compartment algorithm for each participant, as incorporated in the TIVA Trainer software. Propofol infusion commenced with a target effect-site concentration of 0.6 µg/ml and oxygen was titrated to maintain SpO2 above 96%. If Ramsay level was lower than 5, the concentration was slowly increased by increments of 0.3 µg/ml with repeated assessments of responsiveness between increments to obtain a Ramsay score of 5. Once participants stopped responding to verbal commands, were unable to engage in conversation, and were rousable only to physical stimulation they were considered to be at Ramsay level 5. The mean estimated effect-site propofol concentration was 2.48 (1.82-3.14) µg/ml, and the mean estimated plasma propofol concentration was 2.68 (1.92-3.44) µg/ml. Mean total mass of propofol administered was 486.58 (373.30-599.86) mg. The variability of these concentrations and doses is typical for studies of the pharmacokinetics and pharmacodynamics of propofol (SI). For both sessions, prior to the scanning, volunteers were asked to perform a basic verbal recall memory test and a computerized (4 minute) auditory target detection task (SI), which further assessed each individual’s wakefulness/deep anesthesia level independently of the anesthesia team. Scanning commenced only once the agreement among the 3 anesthesia assessors on the Ramsey level 5 was consistent with the lack of response in both verbal and computerized behavioral tests.

Scanning took place in a research not hospital setting, thus, breathing in the deeply anesthetized individuals could not be protected by intubation and was kept under spontaneous individual control. Therefore, although individuals were monitored closely by two anesthesiologists, airway security was at risk during scanning and time inside the scanner was kept at the minimum to ensure return to normal breathing. Thus, safety concerns for the deeply anesthetized individuals dictated that the novel condition of the naturalistic audio story be presented first. The baseline condition of the resting state was considered of secondary importance, as it has been reported previously in deep sedation condition of clinical studies. Therefore, this condition was acquired after the story condition across participants. However, the mean estimated effect-site propofol concentration and the mean estimated plasma propofol concentrations were kept stable by the pharmacokinetic model delivered via the TIVA Trainer infusion pump throughout the deep sedation session, and the lack of significant differences in the frame-wise movement parameters (assessed according to Power et al. (2012))(78) between the story and the resting state conditions further suggested no difference in the level of sedation between the two conditions. For similar safety reasons, data on the meaningless baseline (scrambled version of the audio story) that was designed to clarify processing mechanisms in wakeful individuals, was not collected in deeply anesthetized individuals. Throughout the deep sedation scanning session, the participant’s behavioral profile was monitored inside the scanner room by the anesthesia nurse and one of the anesthesiologists and outside from the scanner control room, with an infrared camera that displayed the participant’s face. No movement, fluctuations of sedation, or any other state change, was observed during the deep sedation scanning for any of the participants included in the study.

### Patients

The severely brain-injured patients were selected based on their clinical diagnoses (i.e., VS/MCS/LIS, at the time of fMRI data acquisition) to form a convenience sample of the disorders of consciousness (DoC) population. No previous fMRI data was available for any of the patients at the time of scanning. Prior to commencing the scanning sessions, all VS/MCS patients were tested behaviorally at their bedside (outside of the scanner) with the Comma Recovery Scale-Revised (CRS-R) (79). At the bedside behavioral testing, six patients met the recognized criteria for the vegetative state (VS), four for the minimally conscious state (MCS), and one for the locked-in syndrome (LIS) (full description of behavioral scores in Table S3). LIS describes an individual who, as a result of acute injury to the brain stem, has (almost) entirely lost the ability to produce motor actions, apart for small, but reproducible eye movements that confirm the presence of consciousness (80). The patients’ demographic and clinical data are summarized in Table S2, S3, and the structural, functional MRI assessment data in Figures 6, S1, S5, S6.

Inside the scanner, each patient underwent a previously established fMRI-based protocol for assessing auditory perception and detecting covert awareness (22, 28) (Figure 6, Figure S5), in the same visit as the audio story scan to help establish the genuine status of consciousness. Prior to assessing command-following, we assessed auditory perception to ensure that it could not have been a limiting factor to producing willful brain responses. Patients had complex underlying medical states, including head flexion and overall muscle rigidity, tracheal tubes for assisted feeding and suctioning, etc., and the highly physically constraining scanning environment compromised their comfort. Some could not lie flat for long periods, others needed frequent suctioning, and other still became agitated after an initial brief period in the scanner. Therefore, to limit patient discomfort, time in the scanner was kept at a minimum and data on the meaningless baseline (scrambled version of the audio story) was not collected.

### fMRI Acquisition and Analysis

#### Healthy individuals

Functional images were acquired on a 3 Tesla Siemens Tim Trio system, with a 32-channel head coil. Standard preprocessing procedures and data analyses were performed with SPM8 and the AA pipeline software (81). In the processing pipeline, a temporal high-pass filter with a cut-off of 1/128 Hz was applied and movement was accounted for by regressing out the 6 motion parameters (x, y, z, roll, pitch, yaw). Additionally, frame-wise movement parameters according to Power et al. (2012) (78) were computed. Prior to analyses, the first five scans of each session were discarded to achieve T1 equilibrium and to allow participants to adjust to the noise of the scanner. To avoid the formation of artificial anti-correlations, a confounding effect previously reported by Murphy and others (82-83), we performed no global signal regression. Group-level correlational analyses explored, for each voxel, the inter-subject correlation in brain activity, by measuring the correlation of each subject’s time-course with the mean time-course of all other subjects. Significant clusters/voxels survived the p<0.05 threshold, corrected for multiple comparisons with the family wise error (FWE). Functional connectivity (FC) was measured by computing via Pearson correlation the similarity of the fMRI time-courses of regions of interest (ROI)—based on well-established landmark ROIs from the resting state literature (84) (Table S1)—*within* and *between* different networks (85) (SI). As this measure of connectivity reflected the degree of similarity between the networks’ functional time-courses, an increase/up-regulation of connectivity indicated more similar time-courses between networks, and thus a loss of functional differentiation. Thus, ‘differentiation’ in this context is measured as the inverse of the Pearson correlation value and must not be confused with measures used in other approaches (48). We note that Pearson correlation is a simple FC measure that, while advantageous for its minimal assumptions regarding the true nature of brain interactions and breath of its use in the neuroscientific literature, does not directly imply causal relations between neural regions. However, it is an adequate measure of FC for our purposes, because the time-course and spatial extent of the auditory and fronto-parietal networks encompassed a vast swath of the hierarchical processing cascade and, thus, many regions of cause-effect space were triggered by the stimulus. Their FC, as measured through Pearson correlation, reflected their interactions over the several minutes and the resulting computations on the information content of the auditory inputs. Future studies will also investigate the connectivity between these regions by using direct measures of causal relationships (86-87). T-tests used to explore effects of interest between functional connectivity and cognitive performance were Bonferroni corrected for multiple comparisons.

#### Severely brain-injured patients

Patient scanning was performed using the same 3 Tesla Siemens Tim Trio system, 32-channel head coil, and data acquisition parameters as for the healthy participants. The same data preprocessing and analyses procedures as for healthy participants were applied to patient data. The patients’ spontaneous arousal during the audio story condition was monitored with an infrared camera placed inside the scanner. One patient (P7) fell asleep in the scanner for the entirely of the session and thus, showed no neuroimaging evidence of awareness despite an MCS diagnosis. The extent of information processing in individual patients (Figure S6) was investigated with a novel technique developed by Naci and colleagues (30, 32). This approach did not involve normalization to a healthy template, nor did it constrain the patient’s expected brain activity based on the localization of the effect in healthy controls. Instead, the time-course of brain activity in healthy controls served to build a strong prediction for the temporal evolution of brain activity in the patients. The precise location of a patient’s brain activity was expected to deviate from that of the healthy controls’. Not only is this naturally the case for individual healthy participants, but also, importantly, it is to be expected in brain-injured patients as a result of structural and concomitant functional re-organization of the brain. Nevertheless, a spatial heuristic based on the controls’ data informed the interpretation of the patients’ results, helping to infer the nature of the underlying residual brain function. In summary, drawing comparisons in the temporal domain enabled direct relation of the healthy controls’ activation to that of brain-injured patients, while avoiding stringent spatial constraints on the patients’ functional anatomy (Figure S6). By contrast, for the analysis of functional connectivity based on a set of network nodes pre-defined in the healthy literature in the MNI standard neurological space, each patient’s brain was normalized to the healthy template. We reasoned that any damage within the regions of interest in each patient’s brain would add noise to the brain activity measurement and reduce the power to detect an effect. Therefore, any results in brain injured patients, that aligned with a-priory hypotheses based on the anesthesia study were highly unlikely given the heterogeneous structural preservation and would present a conservative estimate of the underlying effect.

## Author Contributions

Conceptualization, L.N.; Methodology, L.N., A.H., R.C.; Recruitment, E.H., A.M., M.A., L.G.; Data Acquisition, L.N., E.H., A.M., M.A, S.N., M.A., C.H.; Formal Analyses, L.N., A.H., E.H. R.C.; Writing, L.N.; Supervision, A.M.O.; Funding and resource acquisition, M.A., C.H., A.M.O.

## Acknowledgements

We thank Dr. Conor Wild and Ms. Leah Sinai for their help with the scrambled audio stimulus generation and data acquisition, and Dr. Tim Bayne for his valuable feedback. This research was supported by the L’Oréal-Unesco for Women in Science Excellence Research Fellowship and the Wellcome Trust Institutional Strategic Support Fund to L.N. and a Canada Excellence Research Chair to A.M.O.

## Conflict of Interest

The authors declare no conflict of interest.

## Data Availability Statement

The datasets generated during and/or analysed during the current study are available from the corresponding author on reasonable request.

